# Making ATP fast and slow: do yeasts play a mixed strategy to metabolize glucose?

**DOI:** 10.1101/540757

**Authors:** Hadiseh Safdari, Mehdi Sadeghi, Ata Kalirad

**Affiliations:** School of Biological Science, Institute for Research in Fundamental Sciences (IPM), Tehran, Iran; National Institute of Genetic Engineering and Biotechnology (NIGEB), Tehran, Iran

**Keywords:** metabolic pathways, evolutionary game theory, metabolic regulation

## Abstract

The ability of some microorganisms to switch from respiration to fermentation in the presence of oxygen -the so-called Crabtree effect- has been a fascinating subject of study on theoretical and experimental fronts. Game-theoretical approaches have been routinely used to examine and explain the way a microorganism, such as yeast, would switch between the two ATP-producing pathways, i.e., respiration and fermentation. Here we attempt to explain the switch between respiration and fermentation in yeast by constructing a simple metabolic switch. We then utilize an individual-based model, in which each individual is equipped with all the relevant chemical reactions, to see how cells equipped with such metabolic switch would behave in different conditions. We further investigate our proposed metabolic switch using the game-theoretical approach. Based on this model, we postulate that individuals play a mixed game of glucose metabolism in the population. This approach not only sheds some light in the varieties of metabolic regulations that can be utilized by the individual in the population in competition with others for a common resource, it would also allow a better understanding of the causes of the Warburg effect and similar phenomena observed in nature.

## Introduction

The viability of an organism depends on the fit between its phenotype and the environment it inhabits; an environment that includes the abiotic and biotic factors. When discussing the characteristics of living entities, Waddington enumerated three types of temporal changes that shape a living system: evolution, development, and physiology [1]. The different strategies that a microorganism utilizes to tackle the environmental needs can be explained either in terms of Waddington’s first type of temporal changes (evolution) or the last type of change (physiology).

In an evolutionary explanation, a given genotype have been shaped by evolutionary processes - i.e., natural selection, drift, mutation, & recombination. The genotype would translate into the phenotype which would affect the survival strategy of the organism. The physiological response, on the other hand, should consist of an apparatus that has been shaped by evolution but is capable of responding to the environmental cues in much shorter time spans.

The “decision-making” in *λ* phage is a textbook example of an apparatus - in this case, a genetic switch. The *λ* phage switch has been shaped by natural selection so that it would give the virus a choice between two possible strategies (lysing the host or integrating within the host’s genome), thus equipping the phage to respond to the transient conditions in a timely manner [2]. The “decision-making” approach has been widely used to better understand the intricacies biological phenomena (reviewed in [3]).

The ATP-producing pathways in *Saccharomyces cerevisiae* is a perfect example of a set of strategies that can be explained differently by invoking evolution or physiology. A yeast can either convert glucose through fermentation, a process that is fast but low yields, ≈ 2 moles of ATP per 1 mole of glucose, or go down the slower path -i.e., respiration-, and produce ≈ 32 moles of ATP per 1 mole of glucose [4]. Is the strategies utilized by a yeast a fixed response hardwired in its genome by the natural selection, or is it a physiological response, a distant relative of *λ* phage decision-making apparatus?

The characteristics of fermentation –i.e., fast and low in yield– and respiration –i.e., slow but high in yield– can be easily reformulated using a game-theoretical approach, where fermentation is cast as the “cheater” strategy and the respiration as the “cooperative” one. If the choice of the ATP-producing pathway is a fixed behavior, determined by the genetics, then one could assume that the “selfish” strains would follow the dictum described by Hobbes as “the war of every man against every man” [5] and simply consume the glucose as fast as possible - through fermentation - to outcompete other yeasts in the environment. The game-theoretical approach is an attempt to explain why in nature microorganisms capable of both respiration and fermentation, do not always follow the principle of the maximization of molar yield [6].

Pfeiffer *et al*. [7] led the charge in utilizing the game-theoretical framework to address why yeasts would sometimes choose to ferment and other times to respirate: if every yeast would ferment, then the pool of glucose would drain so fast that each yeast would only get few ATPs. This “tragedy of the commons” is avoidable through respiration, which is viewed through game-theoretical prism as cooperation. Others further analyzed and expanded this approach to explain the trade-off between yield and rate and the dynamic nature of pay-offs for each ATP-producing strategy in yeast (e.g., see [8, 9]). Aside from the trade-off hypothesis, some suggested that the accumulation of ethanol can be utilized by certain strains of yeasts to poison less-alcohol-tolerant strains in their niche [10]. The conditions of cooperation in microorganisms have been investigated experimentally as well (e.g., [8]).

Even the game-theoretical approach described above does not distinguish between a situation where different strains play “selfish” or “cooperative”, something that would require the hand of evolution to intervene, or a scenario in which each yeast has the apparatus to play selfish AND cooperative in different measures to suit its temporal needs. In fact, in reformulating the fermentation/respiration dichotomy in the mould of evolutionary game theory, the “cheaters” and “cooperators” are usually considered different strains that compete for a shared resource (e.g., [11–13]).

In our view, there is no reason to preclude the possibility that individuals do mix these two ATP-producing strategies depending on the environmental conditions. Here, we propose a simple form of such model that allow individual yeast cells to play a mixture of fermentation and respiration. When dealing with a population of organisms showing a mixture of two strategies –e.g., 25% cheating to 75% cooperation – the population-level phenomenon can be caused by two distinct situations at the level of the individuals: a) either 25% of individuals are exclusively utilizing the cheating strategy, while 75% exclusively cooperate, or b) each individual mixes cheating with cooperation in 1 to 3 ratio. The same dilemma is applicable to *S*.*cerevisiae*, or any other microorganism capable of fermentation and respiration.

Here, we attempt to construct a simple regulatory network which makes it possible for a microorganism to combine fermentation and respiration as a mixed strategy. We utilize two models to investigate the ramifications of this regulatory network: firstly, simulating the chemical reactions relevant to the ATP-producing pathways at individual and population levels, and secondly, using a game-theoretical approach. Our results from the population-level modelling of chemical reactions show that such regulatory network affords yeasts to utilize a mixed strategy. The results from the game-theoretical framework demonstrates the inviability of pure strategy in this system. Finally, we suggest an experimental approach to distinguish between a population of yeasts taking advantage of a mixed strategy versus a population that consists of distinct cheaters and cooperators.

## Model

### Metabolic regulatory network for playing a mixed strategy

For yeasts to be able to mix cheating, i.e., glycolysis, with cooperation, i.e., utilizing the TCA cycle, we postulate a regulatory network that is partly supported by the experimental works conducted on *S*.*cerevisiae*. In the ATP-producing pathways, pyruvate kinase seems to play an interesting regulatory role: in yeast, two paralogs of pyruvate kinase (PYK) exist: *PYK1* and *PYK2*. A comprehensive study by Gruning *et al*. [14] concludes that a switch from expressing *PYK1* to *PYK2* corresponds with a shift from fermentation to respiration and the accumulation of PEP suppresses the glycolytic pathway. Boles *et al*. [15] showed that the expression of *PYK2* is suppressed by glucose. Based on these experimental observations, we constructed a regulatory network that modulates the ATP-producing pathways (figure 1). The results from the experimental evolution in yeast is in accordance with our regulatory network: Comparing the gene expression patterns between different populations of *S*.*cerevisiaein* shows a decrease in *PYK1* expression in the evolved lines with reduced ethanol production in comparison with the parental lines [16]. One can suspect that many interactions can be included in this network, but a simple model like this, if correct, may explain the usage of ATP-producing pathways in a cell.

**Fig 1.**
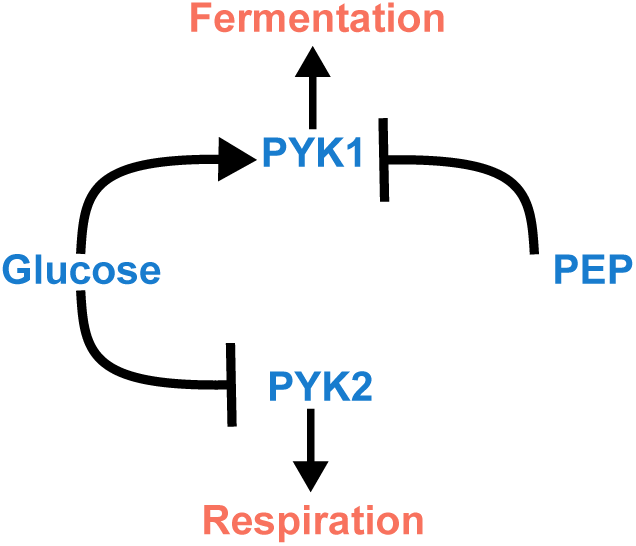
A simple regulatory network to explain utilizing both ATP-producing pathways. Our proposed regulatory network imagines a simple switch between fermentation and respiration based on the experimental data.

### Modelling respiration and fermentation

In modelling glycolysis and the TCA cycle in a digital microorganism, we condensed the myriad of reactions into a few main reactions (Table 1). To simulate the chemical reactions we used Gillespie’s first-reaction method [17]. Since the reaction constants are mesoscopic and dissimilar to deterministic rate constants [18], we choose values that result in biologically-reasonable behavior in our digital microorganism.

**Table 1.** The chemical reactions used to model respiration and fermentation.

| # | Reaction | Propensity ($a_i$) | Parameters |
| --- | --- | --- | --- |
| 1 | $GLC + 2NAD^+ \longrightarrow 2PEP + 2NADH$ | $k_1 \times \#(GLC) \times \#(NAD^+)^2$ | $k_1 = 10^{-6}$ |
| 2 | $PEP \longrightarrow PYR + ATP$ | $k_2 \times \#(PEP)$ | $k_2 = 2$ |
| 3 | $PYR + NADH \xrightarrow{PYK1} EtOH + NAD^+$ | $k_3 \times \#(PYR) \times \#(NADH) \times \#(PYK1)$ | $k_3 = 10^{-2}$ |
| 4 | $EtOH + NAD^+ \longrightarrow PYR + NADH$ | $k_4 \times \#(EtOH) \times \#(NAD^+)$ | $k_4 = 10^{-5}$ |
| 5 | $PYR \xrightarrow{PYK2} 17ATP$ | $k_5 \times \#(PYR) \times \#(PYK2)$ | $k_5 = 10^{-4}$ |
| 6 | $PYK_{1inactive} \longrightarrow PYK_1$ | $\theta_6 \times \#(PYK_{1inactive})$ | See Eq. 1 |
| 7 | $PYK_{2inactive} \longrightarrow PYK_2$ | $k_7 \times \#(PYK_{2inactive})$ | $k_6 = 10^{-3}$ |
| 8 | $PYK_1 \longrightarrow PYK_{1inactive}$ | $\theta_8 \times \#(PYK1)$ | See Eq. 2 |
| 9 | $PYK_2 \longrightarrow PYK_{2inactive}$ | $\theta_9 \times \#(PYK2)$ | See Eq. 3 |

\* GLC: Glucose, PYR: Pyruvate, PEP: Phosphoenolpyruvate, EtOH: Ethanol, PYK: Pyruvate kinase.

In order to factor in the regulatory network (figure 1), in reactions #3 (fermentation) and #5 (the TCA cycle), we multiply the propensities of these reactions by the number of PYK1 and PYK2, respectively. For the activation of PYK1 (reaction #6) a Hill function is used to include the stimulating effect of glucose on PYK1 activation:

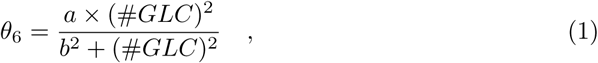

where *a* = 10^−3^ and *b* = 30. The propensity of the PYK1 inactivation reaction (reaction #8) includes a Hill function to take into account the inhibitory effect of PEP on this enzyme:

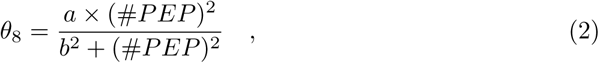

where *a* = 10^−2^ and *b* = 1. PYK2 inactivation is modelled in a similar fashion (reaction #9), where glucose exerts its inhibitory effect on PYK2 by facilitating the PYK2 inactivation reaction through a Hill function:

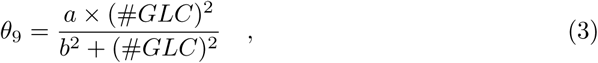

where *a* = 10^−3^ and *b* = 30.

In a population of cells in our model, each cell has its own set of chemical reactions (as described in Table 1). In each step of a simulation, all the first reactions in every cell is determined using Gillespie’s method. Then, the cells execute their first reactions in the order of their reaction times –i.e., the first cell to execute its reaction has the lowest *τ*, while the last cell to proceed with its reaction has the largest *τ*. Afterwards, the population-level clock moves *τ*_max_ forward. During the simulations the population size remains constant.

### Evolutionary Game-theoretical model of mixed metabolic strategy

The occurrence of respiration or fermentation pathways could be treated as a decision making process by rational players. Since they share the same resource for energy, the respiration pathway is considered as a cooperative strategy – with high yield/low rate– and the fermentation as a cheater strategy – with low yield/ high rate. Therefore, at any time, the population consists of a mixture of cooperators and cheaters. The goal in this approach is to find the evolutionary stable strategy (ESS), i.e., a set of strategies chosen by players which cannot be invaded by other strategies. In this approach, the selection criterion between these two choices would be based on the payoff matrix which shows the benefits gained by players as the result of their decisions. The payoffs depend on the amount of glucose as a crucial environmental factor.

The energetic benefit for a player utilizing the respiration pathway is shown by *V* which is much more than the energetic benefit of the fermentation pathway, *W*. However, in each of the metabolic pathways, cells as players have to consume energy to synthesize enzymes, *C* and *D* in respiration and fermentation respectively and *C > D*.

The symmetric payoff matrix of the game (Table 2) represents the gain acquired by player 1 against player 2. According to this matrix, if two players choose the respiration pathway, the payoff –which is the gain in the respiration minus its cost– will be distributed equally between them. On the contrary, if both choose anaerobic pathway, again they should split the payoff (i.e., the gain in the fermentation case mines its enzymatic cost). However, the payoff for a cooperative strategy against a cheating strategy and vice versa, implicitly depends on the amount of glucose, as the rate of respiration and fermentation pathways are highly correlated with the glucose value. We define the ratio of the above rates with *n* as a nonlinear Hill function,

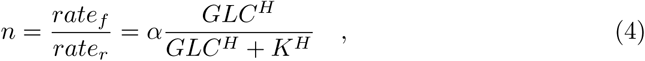

in which *GLC* and *H* indicates the glucose value and Hill exponent and *K* is the half saturation value. This ratio, *n*, affects the off diagonal elements in the the payoff matrix (Table 2). For large amounts of glucose, *n* approaches its maximum value *α*; therefore, the payoff for a cheating strategy against a cooperation would be higher,

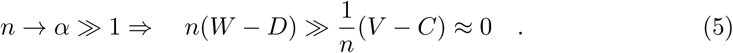

As a result, when glucose is abundant, the fermentation/cheating would be the dominant strategy. On the contrary, in the case of glucose deficiency, *n* approaches zero. In this scenario, the cooperation would be reaping more benefit against cheating and would be the dominant strategy in this situation,

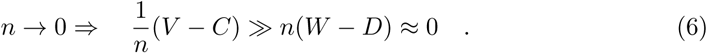

In a medium where the concentrations of nutrients constantly changes, the encounter between cooperation and cheating for a shared resource could be considered as a dynamic Hawk-Dove game. When the glucose is abundant, being a Hawk is the best response, in contrast, a Dove strategy will dominate the population when there is a shortage of glucose. When a moderate amount of glucose exists in the environment, neither of the strategies could invade the other one. In this range of resources, both subpopulations coexist simultaneously.

**Table 2.** The payoff matrix of the game between respiration and fermentation strategies; the elements show the benefits to players 1 and 2 – from left to right, respectively– while the strategies in rows are selected by player 1.

|  |  | player 2 |  |
| --- | --- | --- | --- |
|  |  | respiration | fermentation |
| player 1 | respiration | $(\frac{1}{2}(V - C), \frac{1}{2}(V - C))$ | $(\frac{1}{n}(V - C), n(W - D))$ |
| | fermentation | $(n(W - D), \frac{1}{n}(V - C))$ | $(\frac{1}{2}(W - D), \frac{1}{2}(W - D))$ |

We can consider the dynamic of the strategies over time by the replicator equation. Assuming that *x* fraction of the population choose the strategy *r* with fitness *f*_*r*_, and 1 − *x* the *f* strategy, with fitness *f*_*f*_, the replicator equation would be:

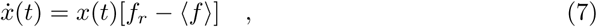

in which ⟨*f*⟩= *f*_*r*_ + *f*_*f*_ is the mean fitness of the population. This deterministic, nonlinear game theoretical equation, by considering the frequency dependent fitness for each strategy, calculates its frequency in the population over time. Based on the payoff matrix *A* in table 2, fitness of the cooperators could be defined as *f*_*r*_ = *Ax*, and ⟨*f*⟩ = *x*^*T*^ *Ax*, where *x*^*T*^ is the transpose of *x*. We can rewrite the Eq. 7 as,

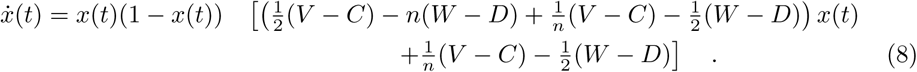

The glucose effect is introduced through a new equation coupled with the replicator equation to reflect the consumption of glucose by cells and its influence on the payoffs:

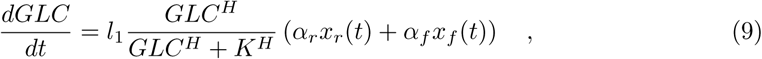

with *K* as the half saturation constant, and *H* the Hill exponent. The *α*_*r*_ and *α*_*f*_, respectively, show the rate of respiration and fermentation pathways.

## Results

### The bias of the regulatory metabolic network in a single cell depends on the amount of glucose

While the raison d’être for the regulatory network espoused in this manuscript is to explain the metabolic behavior of cells at the population level, it has ramifications for a single cell in isolation as well. If a small number of glucose molecules are present in the environment, a cell would “choose” the respiration over fermentation (figure 2A). The low glucose means weaker activation of PYK1 and weaker repression of PYK2. Such behavior is consistent with the yield-rate hypothesis as well, since in the absence of competition, there is no need for rapid consumption of glucose through fermentation (“cheating”) and the payoff for respiration will be paramount.

**Fig 2.**
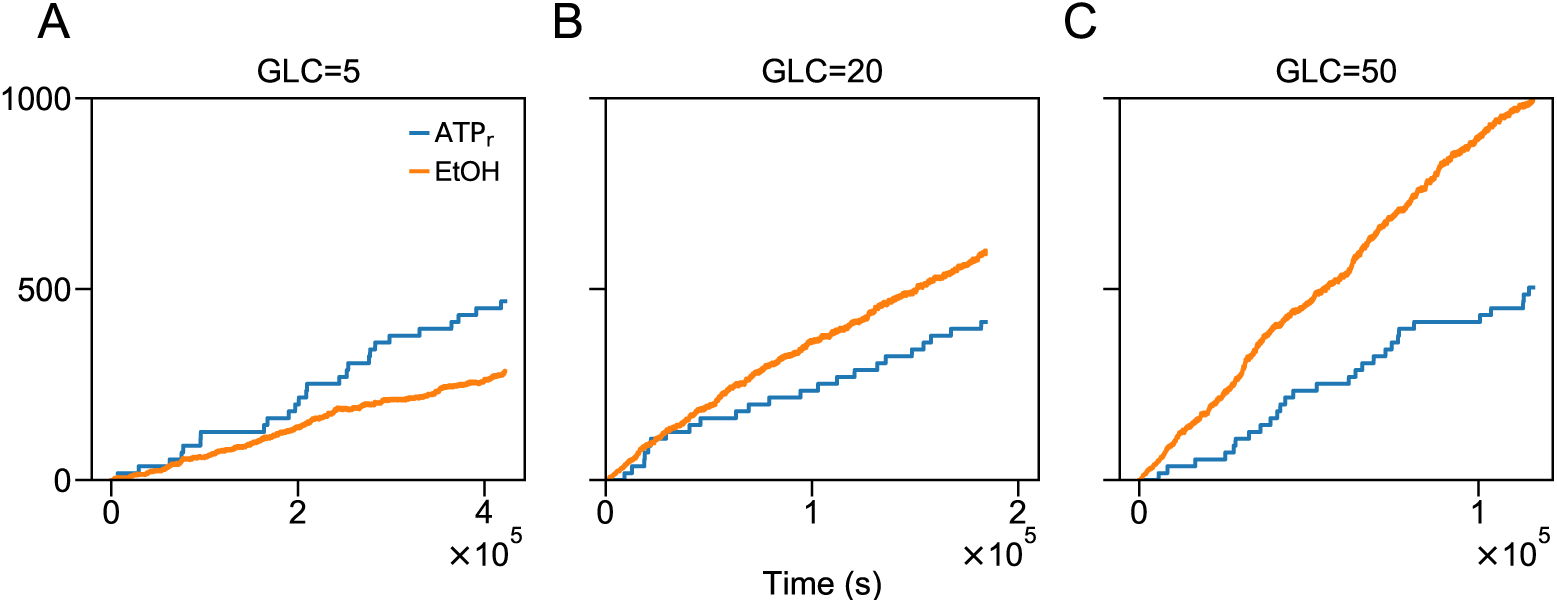
The effect of glucose concentration on the metabolic regulation of a single yeast cell. Result based on the behavior of a single digital yeast. The number of available glucose during simulation remained constant.

Since glucose is the key regulatory element in our network, increasing the number of glucose molecules drives the regulatory networks towards fermentation (figure 2B and 2C). When the energy source in the environment is more abundant, the balance of the metabolic scale tilts towards faster energy production –i.e., fermentation– since there is no benefit to preferring the slower but more efficient path. It is worth-noting again that in our model, a cell, even in isolation, plays a “mixed” strategy and the choice between higher yield or faster rate is never binary.

### The usage of ATP-producing pathways in a homogenous population depend on the population size

Interestingly, in the chemical-reactions model, the cells equipped with the regulatory metabolic network behave as one would expect within the confines of the yield-rate hypothesis. In the extreme case, where there are only two cells in a homogenous environment –i.e., an environment lacking any spatial structure– and only 2 molecules of glucose are accessible to them, the cells would opt for the “cooperative” strategy (respiration), since no “cheating” is justifiable with such a limited resource (figure 3A). Increasing the amount of accessible glucose to 100 molecules, immediately affect the behavior of two cells: now they utilize a mixed strategy of respiration/fermentation where fermentation is dominant (figure 3B).

**Fig 3.**
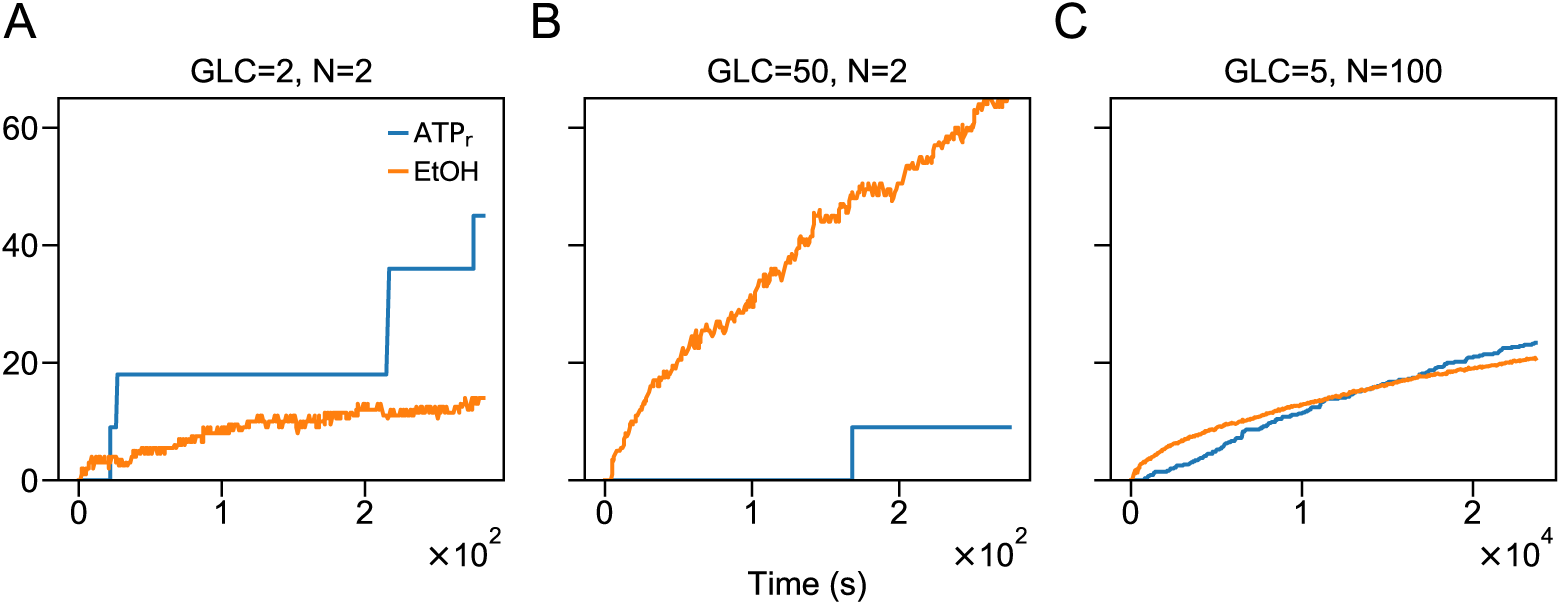
The population size differentially affects the metabolic regulation of a single yeast cell at different glucose concentrations. The number of available glucose during simulation remained constant.

Increasing the number of cells to 100, with a low number of accessible glucose (5), results in a mixed strategy of respiration/fermentation. The smaller proportion of the average number of accessible glucose per cell, opting for higher yield is once again the priority (figure 3C).

### The replicator dynamics indicates no pure strategy

Figure (4) shows the amount of glucose over time; as well, the changes in the frequency of two strategies. As it can be seen, for high value of glucose, almost all members of the population follow the cheater strategy as it has the highest payoff and does better against the other strategy, i.e. it is the ESS; however, by glucose consumption and decrease in its level, none of the strategies could invade the other one, therefore, there is not a pure strategy as the ESS. Under this circumstance, the ESS would be a mixture of the cheating and cooperation. For small amounts of glucose, the cooperative strategy has the highest payoff. As a result, it would be the dominant strategy which plays better than cheating; hence, all the population members have the tendency to this choice.

## Discussion

The importance of the fermentation/respiration dichotomy in microorganisms is not a mere theoretical curiosity; the well-known tendency of cancerous cells to consume large amounts of glucose and metabolize it via fermentation, while oxygen is present in their *umwelt*, has been extensively studied since Ott Warburg observed it [19]. In spite of these studies, the biological causes of the Warburg effect have remained largely unresolved [20]. Understanding the underlying biology of the Crabtree effect -i.e., the ability of microorganism to produce ethanol in the presence of oxygen is of extreme importance in the industrial endeavor to optimize ethanol production in yeast (reviewed in [21]).

Studying the alternative metabolic strategies have been a rather fruitfulenterprise. Generally, different strategies are treated as distinct “strain” that do compete for a shared resource (e.g., see [22]). This approach simplifies the problem and allows to treat the problem similar to studying the invasion and fixation of mutant strain in well-mixed and structured populations. Here we propose that, alternatively, there could be no distinct cheaters and cooperators when it comes to alternative ATP-producing pathways.

Our result can provide a molecular basis for further optimization of yeasts for ethanol production. Hitherto, environmental variables such as temperature or pH have been the focus of optimization efforts to increase ethanol production (e.g., [23]). The manipulation of environmental conditions is a coarse-grain method based on the general effect of these conditions on the rate of chemical reactions. Our regulatory network would provide two alternative approaches: at a physiological level one can tune the concentrations of PEP and glucose so that the metabolic switch would tilt towards fermentation, or a site-directed mutagenesis targeting *PYK1* and/or *PYK2* could bias the switch such that the production of ethanol becomes an inevitability.

The notion that cells might combine different strategies is not novel, but our results show that in the case of a microorganism equipped with alternative pathways for metabolizing glucose, a simple experiment can distinguish between a population of microorganisms playing mixed strategy and a population composed of distinct respirators and fermentors. If we sample from a population where each individual respirates or ferments and measure the amount ethanol produced in this sample, or result would vary form those of the population since by chance we could sample many more fermentors (or respirators) and thus proportion of respirators to fermentors would be different in our sample. On the contrary, if we sample from a population of mixed players, we would expect similar results from our sample compared to the population, since the proportion of respirators to fermentors observed at the population-level does not correspond to different individuals utilizing different strategies, but to individuals taking advantage of both pathways (figure 5).

**Fig 4.**
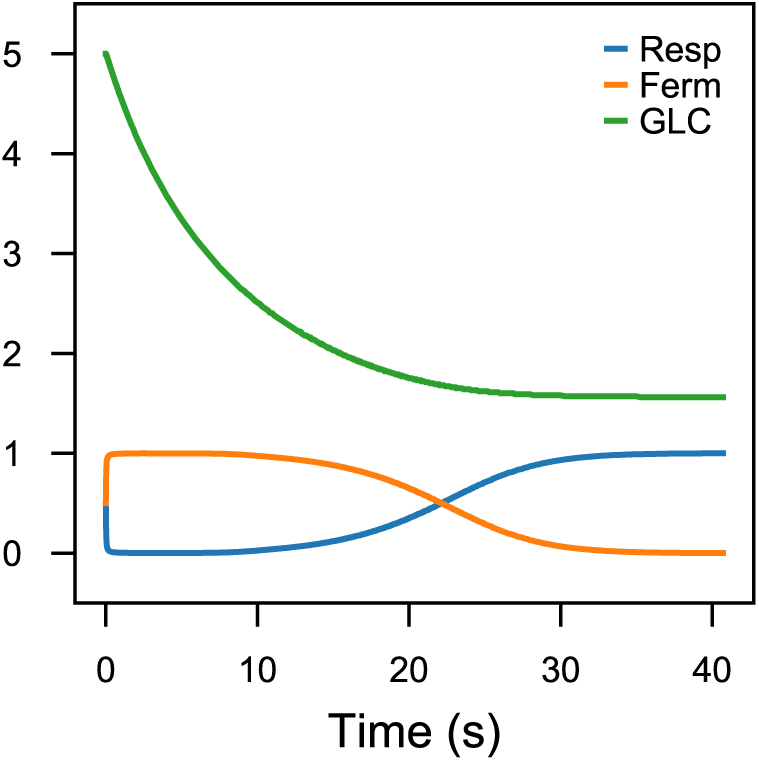
The frequency of two strains population over time as a function of glucose level based on the replicator equation 8 (*α* = 100, *H* = 2, & *K* = 5) and the equation 9 (*l*_1_ = −2, *α* = 100, *H* = 2, & *K* = 100). By reduction in the glucose level, cells with the respiration strategy (cooperators) invade the whole population.

**Fig 5.**
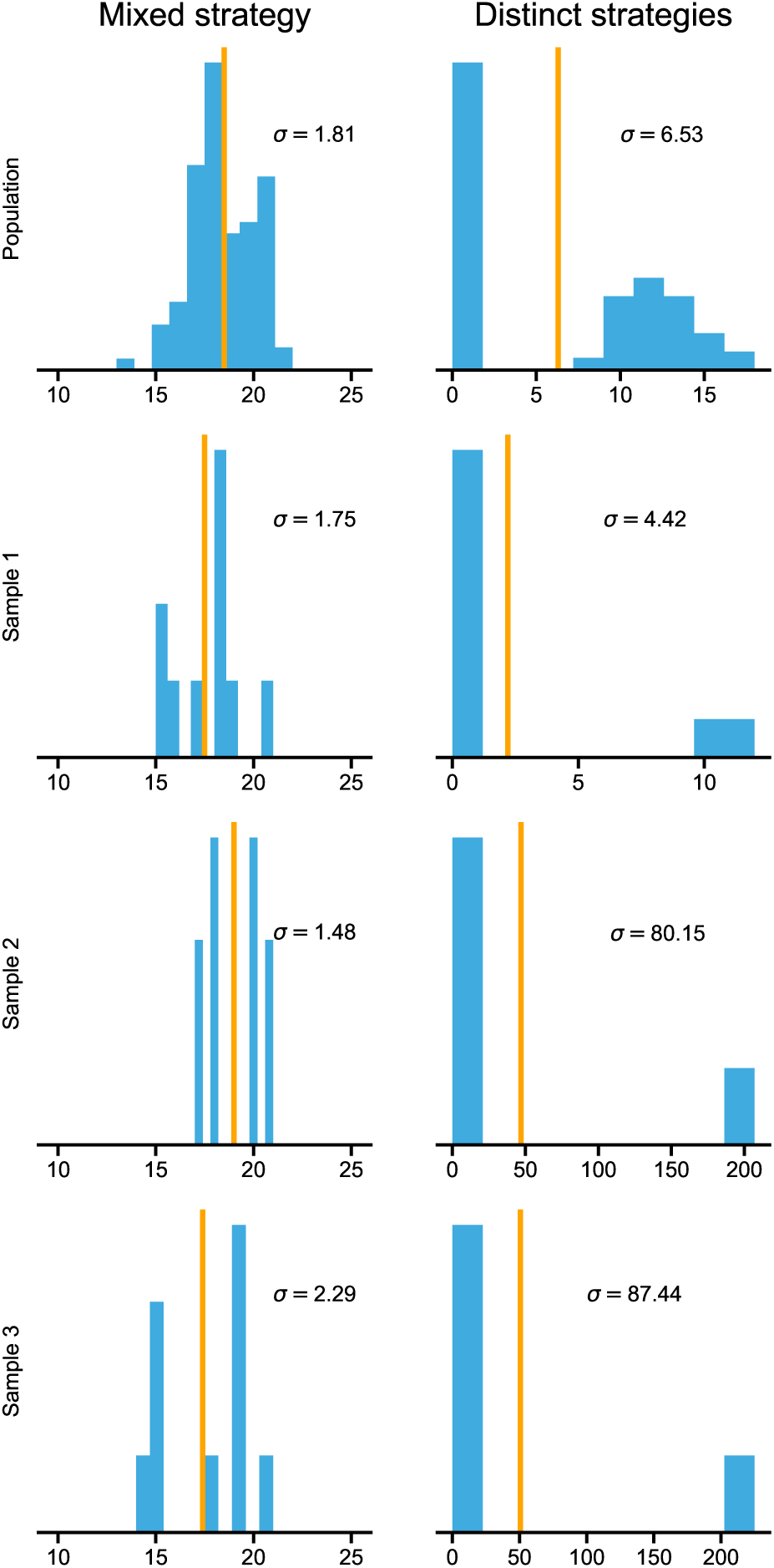
Sampling from the population can distinguish between a population composed of individuals that play a mixed strategy and a population composed of a mixture of strains who each utilize a different strategy. In the mixed strategy case (left column), a population of 100 that utilized a mixed strategy to metabolize glucose are allowed for 100 steps, then sample of size 10 is randomly drawn from this population is allowed to metabolize for 100 Gillespie’s steps. The histograms for the number of ethanols produced by cells at the end of each runs do not show significant variation in variance between the population and the samples, but in a population of yeast who exclusively utilize respiration or fermentation, the dispersion in ethanol production is quite different from a sample to sample. For a population of distinct strategies, the population consisted of equal proportions of respirators and fermentors; this cells utilized the same reactions as table 1, except for *k*_6_ = 10^−1^, *k*_7_ = 10^−5^, *k*_8_ = 0, *k*_9_ = 0 for obligate fermentors and *k*_6_ = 0, *k*_7_ = 0, *k*_8_ = 10_−3_, *k*_9_ = 10_−5_ for obligate respirators; in order for ATP and ethanol productions to become stable at the end of our chemical simulations, populations starts with a shared pool of 1000 glucose molecules and glucose does not replenish during a simulation. Sample populations resume their chemical simulation with a fresh pool of 100 glucose.

It has not escaped our notice that one significant advantage of conceiving the biological details of the metabolic network is the ability to test the veracity of such regulatory scheme. The veracity of our proposed regulatory network can have significant ramification in developing anti-angiogenic drugs, given the over-reliance of cancer cells on glycolysis in place of the TCA cycle [24]. In spite of years of research on the Warburg effect, the biology of this phenomenon is still far from crystalline [20]. Unraveling the changes in the metabolic landscape of a cell as a result of cancer is an endeavor that has only recently been kickstarted [25]. Thus, little is known about the specific roles different metabolites and reactions play in driving cancerous cells to anaerobic ATP production, but our model can be an starting point to explore potential targets to suppress anaerobic ATP production in malignant cells.

## Acknowledgments

We would like to thank Amir Banaei-Esfahani for his early works relevant to this study.

## Availability of data and material

The software used to run all simulations was Python 2.7 and the scripts are available at https://github.com/Kalirad/Making_ATP_fast_and_slow.

## Author contribution

M.S. conceived the model and helped draft the manuscript. H.S. and A.K. performed the simulations, developed the analytical calculations. A.K. drafted the manuscript.

H.S. helped draft the manuscript. M.S. and H.S. critically revised the manuscript. All authors gave final approval for publication.

## Ethics approval and consent to participate

Not applicable.

## Consent for publication

Not applicable.

## References

1. Waddington CH. The Strategy of the Genes: A Discussion of Some Aspects of Theoretical Biology. New York: George Allen & Unwin Ltd; 1957.

2. Shea MA, Ackers GK. The OR control system of bacteriophage lambda. Journal of Molecular Biology. 1985;181(2):211–230. doi:http://dx.doi.org/10.1016/0022-2836(85)90086-5.

3. Perkins TJ, Swain PS. Strategies for cellular decision-making. Molecular Systems Biology. 2009;5(1). doi:10.1038/msb.2009.83.

4. Waddell TG, Repovic P, Melendez-Hevia E, Heinrich R, Montero F. Optimization of glycolysis: A new look at the efficiency of energy coupling. Biochemical Education. 1997;25(4):204–205. doi:10.1016/S0307-4412(97)00131-3.

5. Hobbes T. Leviathan or The Matter, Forme and Power of a Common Wealth Ecclesiasticall and Civil. London: Andrew Crooke; 1651.

6. Schuster S, Pfeiffer T, Fell DA. Is maximization of molar yield in metabolic networks favoured by evolution? Journal of Theoretical Biology. 2008;252(3):497–504. doi:https://doi.org/10.1016/j.jtbi.2007.12.008.

7. Pfeiffer T, Schuster S, Bonhoeffer S. Cooperation and Competition in the Evolution of ATP-Producing Pathways. Science. 2001;292(5516):504–507. doi:10.1126/science.1058079.

8. MacLean RC, Gudelj I. Resource competition and social conflict in experimental populations of yeast. Nature. 2006;441:498–501. doi:https://doi.org/10.1038/nature04624.

9. Frick T, Schuster S. An example of the prisoner’s dilemma in biochemistry. Naturwissenschaften. 2003;90(7):327–331. doi:10.1007/s00114-003-0434-3.

10. Rozpędowska Ez, Hellborg L, Ishchuk OP, Orhan F, Galafassi S, Merico A, et al. Parallel evolution of the make–accumulate–consume strategy in *Saccharomyces* and *Dekkera* yeasts. Nature Communications. 2011;2:302. doi:http://dx.doi.org/10.1038/ncomms1305.

11. Pfeiffer T, Schuster S. Game-theoretical approaches to studying the evolution of biochemical systems. Trends in Biochemical Sciences. 2005;30(1):20–25. doi:https://doi.org/10.1016/j.tibs.2004.11.006.

12. Schuster S, Kreft JU, Schroeter A, Pfeiffer T. Use of Game-Theoretical Methods in Biochemistry and Biophysics. Journal of Biological Physics. 2008;34(1):1–17. doi:10.1007/s10867-008-9101-4.

13. Schuster S, de Figueiredo LF, Schroeter A, Kaleta C. Combining Metabolic Pathway Analysis with Evolutionary Game Theory. Explaining the occurrence of low-yield pathways by an analytic optimization approach. Biosystems. 2011;105(2):147–153. doi:https://doi.org/10.1016/j.biosystems.2011.05.007.

14. Grüning NM, Rinnerthaler M, Bluemlein K, Mülleder M, Wamelink MMC, Lehrach H, et al. Pyruvate Kinase Triggers a Metabolic Feedback Loop that Controls Redox Metabolism in Respiring Cells. Cell Metabolism. 2011;14(3):415–427. doi:10.1016/j.cmet.2011.06.017.

15. Boles E, Schulte F, Miosga T, Freidel K, Schlüter E, Zimmermann FK, et al. Characterization of a glucose-repressed pyruvate kinase (Pyk2p) in Saccharomyces cerevisiae that is catalytically insensitive to fructose-1,6-bisphosphate. Journal of Bacteriology. 1997;179(9):2987–2993. doi:10.1128/jb.179.9.2987-2993.1997.

16. Ferea TL, Botstein D, Brown PO, Rosenzweig RF. Systematic changes in gene expression patterns following adaptive evolution in yeast. Proceedings of the National Academy of Sciences. 1999;96(17):9721–9726. doi:10.1073/pnas.96.17.9721.

17. Gillespie DT. Stochastic Simulation of Chemical Kinetics. Annual Review of Physical Chemistry. 2007;58(1):35–55. doi:10.1146/annurev.physchem.58.032806.104637.

18. Gibson MA, Bruck J. Efficient Exact Stochastic Simulation of Chemical Systems with Many Species and Many Channels. The Journal of Physical Chemistry A. 2000;104(9):1876–1889. doi:10.1021/jp993732q.

19. Warburg O. The Metabolism of Carcinoma Cells. The Journal of Cancer Research. 1925;9(1):148–163. doi:10.1158/jcr.1925.148.

20. Liberti MV, Locasale JW. The Warburg Effect: How Does it Benefit Cancer Cells? Trends in Biochemical Sciences. 2016;41(3):211–218. doi:10.1016/j.tibs.2015.12.001.

21. Cardona CA, Sánchez Ó J. Fuel ethanol production: Process design trends and integration opportunities. Bioresource Technology. 2007;98(12):2415–2457. doi:https://doi.org/10.1016/j.biortech.2007.01.002.

22. Amado A, Fernández L, Huang W, Ferreira FF, Campos PRA. Competing metabolic strategies in a multilevel selection model. Royal Society open science. 2016;3(11):160544; 160544–160544. doi:10.1098/rsos.160544.

23. Arora R, Behera S, Sharma N, Kumar S. A new search for thermotolerant yeasts, its characterization and optimization using response surface methodology for ethanol production. Frontiers in Microbiology. 2015;6:889. doi:10.3389/fmicb.2015.00889.

24. Fitzgerald G, Soro-Arnaiz I, Bock KD. The Warburg Effect in Endothelial Cells and its Potential as an Anti-angiogenic Target in Cancer. Frontiers in Cell and Developmental Biology. 2018;6:100. doi:10.3389/fcell.2018.00100.

25. Jayaraman A, Kumar P, Marin S, de Atauri P, Mateo F, M Thomson T, et al. Untargeted metabolomics reveals distinct metabolic reprogramming in endothelial cells co-cultured with CSC and non-CSC prostate cancer cell subpopulations. PLOS ONE. 2018;13(2):e0192175–.

